# Functional fractionation of large-scale brain networks in the human subcortex

**DOI:** 10.64898/2025.12.23.696301

**Authors:** Jian Li, Alexander S. Atalay, Mark Olchanyi, Morgan K. Cambareri, Satrajit S. Ghosh, Andreas Horn, Laura Lewis, Emery N. Brown, Bruce Fischl, Hannah C. Kinney, Brian L. Edlow

## Abstract

Brain network mapping plays a crucial role in advancing our understanding of human brain organization and the neuroanatomic foundations of cognition. Historically, the identification of large-scale brain networks has focused on the cerebral cortex. Some cortical networks have been fractionated into subnetworks, yielding valuable insights into their domain-specific cognitive functions. In contrast, functional mapping of large-scale brain networks within subcortical regions remains an emerging and challenging field, hindered by a low signal-to-noise ratio in subcortical functional MRI data and an inability to distinguish networks with substantial spatiotemporal overlap. In this study, we fractionated and identified fifteen spatially overlapped and temporally correlated subnetworks, which can be categorized into four large-scale brain networks. The subcortical functional connectivity patterns of these subnetworks exhibited distinct, yet overlapping, spatial configurations, with widely connected hub nodes identified in the caudate, putamen, hippocampus, and thalamus. These subnetworks are highly reproducible across healthy human brains and provide normative functional atlases, which we release here as a resource for the academic community. As a proof-of-principle demonstration of how the atlases can be used to elucidate the pathophysiology of neuropsychiatric disorders, we show that the spatial patterns of the subnetworks predict the level of consciousness in patients with severe traumatic brain injury. These observations indicate a highly conserved and spatially overlapped subcortical functional architecture in the human brain, providing opportunities to elucidate pathophysiologic mechanisms and develop new neuromodulatory therapies for a broad spectrum of neuropsychiatric diseases.

## Introduction

Brain network mapping is essential for advancing knowledge about human brain organization and the neuroanatomic basis of cognition. Historically, identification of large-scale brain networks has focused on the cerebral cortex^1–3^. Functional neuroimaging techniques, such as functional magnetic resonance imaging (fMRI), have revealed highly reproducible network architectures across the cerebral cortex. With the advent of widely available open-source large fMRI datasets, some large-scale cortical networks have been fractionated or “broken down” into subnetworks^4^ – highly interconnected subsystems that perform distinct functions – yielding insights into network components with domain-specific cognitive correlates. These insights have provided the foundation for development of novel therapeutic approaches in a broad range of neuropsychiatric disorders^5^, which target specific cortical components of canonical brain networks^6^.

In contrast, functional mapping of large-scale brain networks within subcortical regions is an emerging field that has been hindered by methodological barriers. Few studies have attempted to map the subcortical connectivity of large-scale networks in the human brain^7–9^, and fractionation of these networks has not been performed at the subcortical level. Subcortical components of human brain networks play an essential role in the modulation of consciousness^10,11^, homeostasis^12^, and emotion^13^, while also contributing to higher-order cognitive functions^14^. The clinical motivation for subcortical fractionation of large-scale brain networks is to identify and refine therapeutic targets^15,16^, as many pharmacologic^17^, electrical^18^, and ultrasound-based^19^ neuromodulatory therapies aim to stimulate widely connected subcortical hub nodes^5,9^. Functional fractionation of brain networks into subnetworks for the entire brain, including the subcortex, thus has the potential to create new therapeutic opportunities across a broad spectrum of neuropsychiatric disorders.

The most critical methodological barrier to functional fractionation of brain networks, particularly their subcortical components, is the overlapping and correlated nature of these brain networks. Spatial overlap between large-scale brain networks has been widely observed in fMRI data at the cerebral cortical level^20–22^ – a key reason that nomenclature for large-scale brain networks remains controversial^23^. Network overlap is even more prominent at the subcortical level due to the so-called “funnel effect” of cortico-subcortical electrical signaling^24^, whereby neuroanatomically distributed information processing in the cerebral cortex is compressed into neuroanatomically localized processing in smaller subcortical structures^25,26^. For example, the thalamus, long considered a relay station in sensory processing within the sensorimotor network (SMN), is now known to also modulate the default mode network (DMN), which is involved in self-referential thought and mind-wandering^7,27^. Similarly, the basal ganglia – including the putamen, globus pallidus, and subthalamic nucleus – modulate the SMN for motor control, as well as the executive control network (ECN) for higher-order cognitive functions such as attention and decision-making^28^. Furthermore, subcortical regions like the amygdala and ventral striatum are functionally connected with both the salience network (SAL) and ECN, suggesting that these regions mediate a dynamic interplay between emotion, cognition, and sensation^29^. However, traditional fMRI analytic methods, such as seed-based correlation analysis (SCA) and independent component analysis (ICA), are unable to disentangle networks that are overlapped or correlated with each other, because the former assumes a linear and direct point-wise relationship between the seed and the target region^30^, while the latter assumes statistical independence among networks^31–33^.

Another critical challenge in subcortical functional brain mapping is the low signal-to-noise ratio (SNR) in fMRI data, particularly in the subcortex. Subcortical regions exhibit lower blood oxygenation level-dependent (BOLD) signal relative to cortical regions, partially due to differences in vascularization and hemodynamic responses in deep brain areas^34^. This reduction in BOLD signal makes it difficult to reliably detect functional activity in subcortical areas. In addition, subcortical structures are often influenced by physiological “noise”, such as respiration, heart rate, and eye movement, which are more pronounced in deeper brain regions^35^. This physiological noise can obscure neural signals and further reduce the effective SNR^36,37^. Due to these barriers, no “ground truth” or consensus has been established in subcortical functional brain mapping, a gap in knowledge that has been a fundamental barrier to identifying subcortical functional connectivity differences in individuals with neurological or psychiatric disorders.

In this resting-state fMRI (rsfMRI) brain mapping study, we fractionated and identified 15 subnetworks within four large-scale brain networks – the DMN, SMN, visual network (VN), and higher-order cognitive (HOC) network comprised of the ECN, SAL, dorsal and ventral attentional networks (DAN/VAN) – using the Nadam-Accelerated SCAlable and Robust (NASCAR) canonical polyadic decomposition method^7,38,39^. We specifically designed the NASCAR method to separate overlapping brain networks from each other, enabling us to address the methodological challenges that have historically limited subcortical brain mapping. By applying NASCAR to rsfMRI data from 1,000 subjects in the Human Connectome Project (HCP), in combination with bootstrapping and embedding techniques, we revealed the subcortical functional connectivity of large-scale brain networks and their subnetworks, identified their sites of neuroanatomic overlap, and demonstrated that these subnetworks are highly reproducible across the healthy human population. These subnetworks reliably identified from 1,000 HCP subjects provide gold-standard, normative functional atlases for the healthy population. In a proof-of-principle application of these functional atlases, we demonstrate that the spatial pattern of the subnetworks predicts the level of consciousness of individual patients with severe traumatic brain injury (TBI). We release the subcortical functional atlases on OpenNeuro and Lead-DBS, and we provide an interactive online viewing platform for the academic community.

## Results

We present four large-scale brain networks, namly the DMN, SMN, VN, and HOC, along with their fractionated subnetworks. We describe their key nodes within the subcortical regions. Tables S1-S4 provide a comprehensive list of subnetwork activity levels in subcortical structures and sub-nuclei. Table S5 includes the acronyms and abbreviations for anatomical structures referenced in the subsequent sections.

### Default mode network (DMN)

The overall DMN, as shown in Fig. 1, includes PCC, (vm, dm, dl)-PFC, IPL, MTG, and PHG cortically, consistent with prior reports^27,40^. Subcortically, the DMN involves the caudate, anterior, dorsal, medial, and posterior thalamus, hippocampus, amygdala, caudal midbrain, and pons. In contrast to prior reports of DMN fractionation that identified two subnetworks^4^, we identified three subnetworks: 1) the medial temporal lobe (DMN-MTL) subnetwork; 2) the dorsal medial prefrontal cortex (DMN-dmPFC) subnetwork; and 3) a newly described ventral posterior cingulate cortex (DMN-vPCC) subnetwork.

**Fig. 1.**
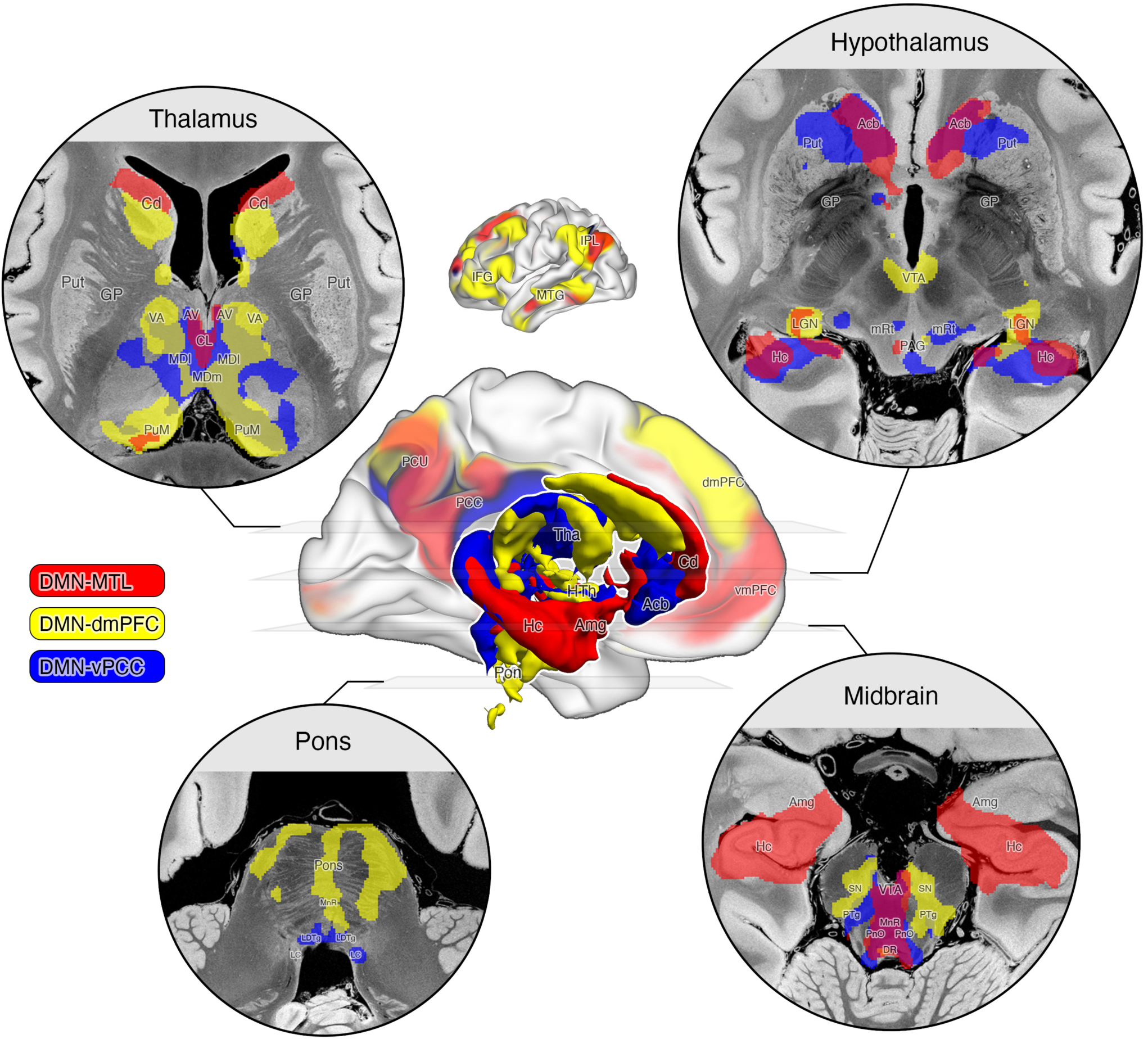
Default mode network (DMN) subnetworks. The medial temporal lobe (DMN-MTL) subnetwork is shown in red, the dorsal medial prefrontal cortex (DMN-dmPFC) subnetwork is shown in yellow, and the ventral posterior cingulate cortex (DMN-vPCC) subnetwork is shown in blue. The medial view of the cortical map of the left hemisphere with the 3D rendering of the subcortical DMN subnetworks are shown in the center. Axial slices through the thalamus, hypothalamus and basal forebrain, midbrain, and the pons are shown in each of the four corners. The lateral view of the cortical map of the left hemisphere is shown on top of the medial view. Refer to the abbreviation section for the annotations of the anatomical referneces.

#### The medial temporal lobe (DMN-MTL) subnetwork (red)

Cortical nodes of DMN-MTL are dPCC, ICG, PCu, PHG, vmPFC medially, and pAG, pSFG, aMTG, pITG laterally. Most of the cortical nodes in this subnetwork were reported by Andrews-Hanna et al.^4^. Although several lateral nodes were not identified there, we retain the naming convention DMN-MTL^4^ based on the extensive medial temporal connectivity in this subnetwork.

Subcortical nodes of DMN-MTL are also positioned medially relative to the other two subnetworks. The anterior hippocampus, amygdala, and the anterior/dorsal caudate are highly involved and unique to DMN-MTL within the DMN. Other subcortical nodes include medial thalamic nuclei such as CL, MDm, PuM, medial brainstem nuclei such as DR, MnR, ventral medial VTA, LDTg, PnO, and hypothalamic nuclei such as AHA, AN, BNST, LH, MM, SO, TM, and VMH.

#### The dorsal medial prefrontal cortex (DMN-dmPFC) subnetwork (yellow)

Cortical nodes of DMN-dmPFC are dmPFC, PCu, dPCC, sps medially, and aAG, the middle section of MTG, TP, dlPFC, and IFG laterally. This subnetwork was identified by Andrews-Hanna and colleagues^4^ without nodes in the lateral frontal regions. We also apply the previously proposed name based on its unique cortical node in dmPFC.

In the subcortical region, strong involvement is observed in the medial/dorsal caudate (body and tail), anterior (AV, VLa, VLp), medial (CL, MDl, MDm), dorsal (LD, LP), and posterior (PuA, PuM, L-Sg, LGN) nuclei of the thalamus, dorsal VTA, PTg, and the pons in the brainstem, and SN in the hypothalamus.

#### The ventral posterior cingulate cortex (DMN-vPCC) subnetwork (blue)

Cortical nodes of DMN-vPCC are vPCC (dorsal to cas), pPCu (rostral to pos), sps medially, and dpAG laterally. For its unique presence in vPCC, we name this subnetwork DMN-vPCC.

Subcortically, the caudate, accumbens, putamen, and hippocampus are highly involved, and the involvement of the anterior putamen is unique to this subnetwork within the DMN. Similar to DMN-dmPFC, DMN-vPCC also includes the anterior (AV), medial (CL, MDl, MDm), dorsal (LD, LP) and pulvinar (PuA, PuM) nuclei in the thalamus. In contrast to DMN-MTL and DMN-dmPFC, this subnetwork has unique and strong involvement in dorsal lateral nuclei of the brainstem such as LC, LDTg, PnO, mRt in addition to medial nuclei such as DR, MnR, and ventral medial VTA. In the hypothalamus, LH, MM, and SN are the key nodes.

### The sensorimotor networks (SMN)

The overall SMN, as shown in Fig. 2, spans the precentral, postcentral gyri, and the paracentral lobule cortically. The SMN is involved in many subcortical structures, with extensive overlap in the thalamus across its three subnetworks.

**Fig. 2.**
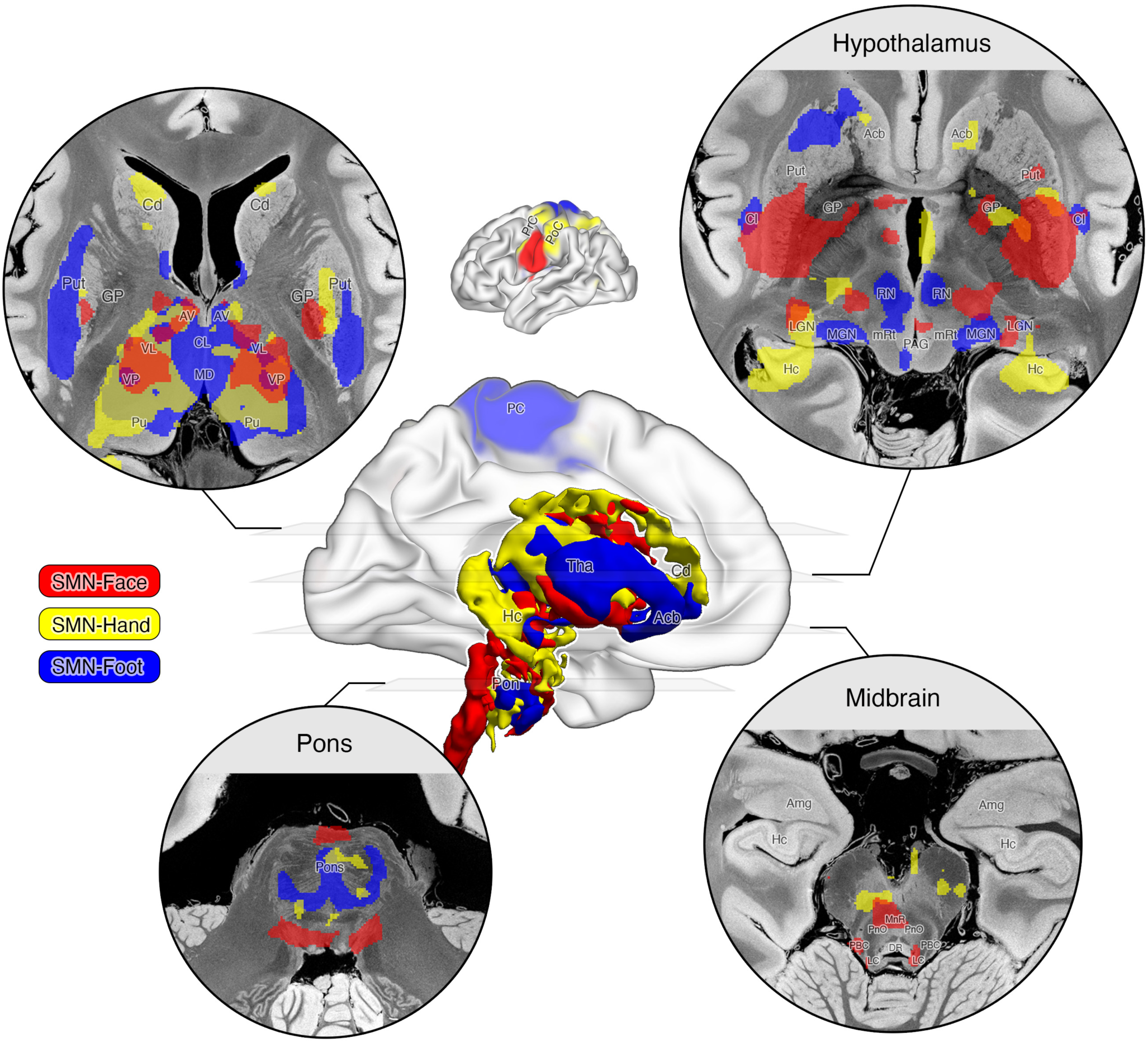
Sensorimotor network (SMN). The face (SMN-Face) subnetwork is shown in red, the hand (SMN-Hand) subnetwork is shown in yellow, and the foot (SMN-Foot) subnetwork is shown in blue. Refer to Fig. 1 for other notations.

#### The face (SMN-Face) subnetwork (red)

The cortical SMN-Face subnetwork involves the inferior PrC and PoC, which are well-established regions for sensorimotor functions of the face/tongue. Unique subcortical nodes of SMN-Face include the superior caudate, ventral posterior putamen, globus pallidus, and claustrum. This subnetwork also involves anterior (AV), dorsal (LD, LP), medial (CL, MDm, MDl), ventral (VA, VLa, VLp, VPL), and posterior (LGN) thalamic nuclei but not pulvinar. In the brainstem, SMN-Face is active in the dorsal lateral pontine tegmentum (LC, PBC). Minor involvement in NBM is observed in the basal forebrain.

#### The hand (SMN-Hand) subnetwork (yellow)

The cortical SMN-Hand subnetwork involves the superior PrC and PoC hand regions. Among the SMN subnetworks, SMN-Hand has unique involvement in the dorsal anterior caudate and posterior hippocampus. SMN-Hand is also involved in the middle section of the putamen. Most thalamic nuclei participate in this subnetwork except for the rostal sections of the thalamus. Brainstem and hypothalamic nuclei are minimally involved in SMN-Hand, except in the basis pontis.

#### The foot (SMN-Foot) subnetwork (blue)

The cortical SMN-Foot subnetwork involves the Pc foot regions. SMN-Foot has a strong involvement in the lateral section of the putamen and partially in sections of the caudate and claustrum that are adjacent to the active lateral putamen areas. Similar to SMN-Hand, SMN-Foot is involved in almost all thalamic nuclei except for the dorsal regions (LD, LP). Involvement in RN, PAG, mRt, and basis pontis is observed in the brainstem. None of the hypothalamic nuclei is highly involved in this subnetwork.

### The visual networks (VN)

The overall VN, as shown in Fig. 3, occupies the visual cortex in the occipital lobe and visual association areas near the IPS and motor cortex. Subcortically, the VN system strongly involves the thalamus, caudate, putamen, accumbens, hippocampus, amygdala, and brainstem, and sparsely the hypothalamus, but not the globus pallidus.

**Fig. 3.**
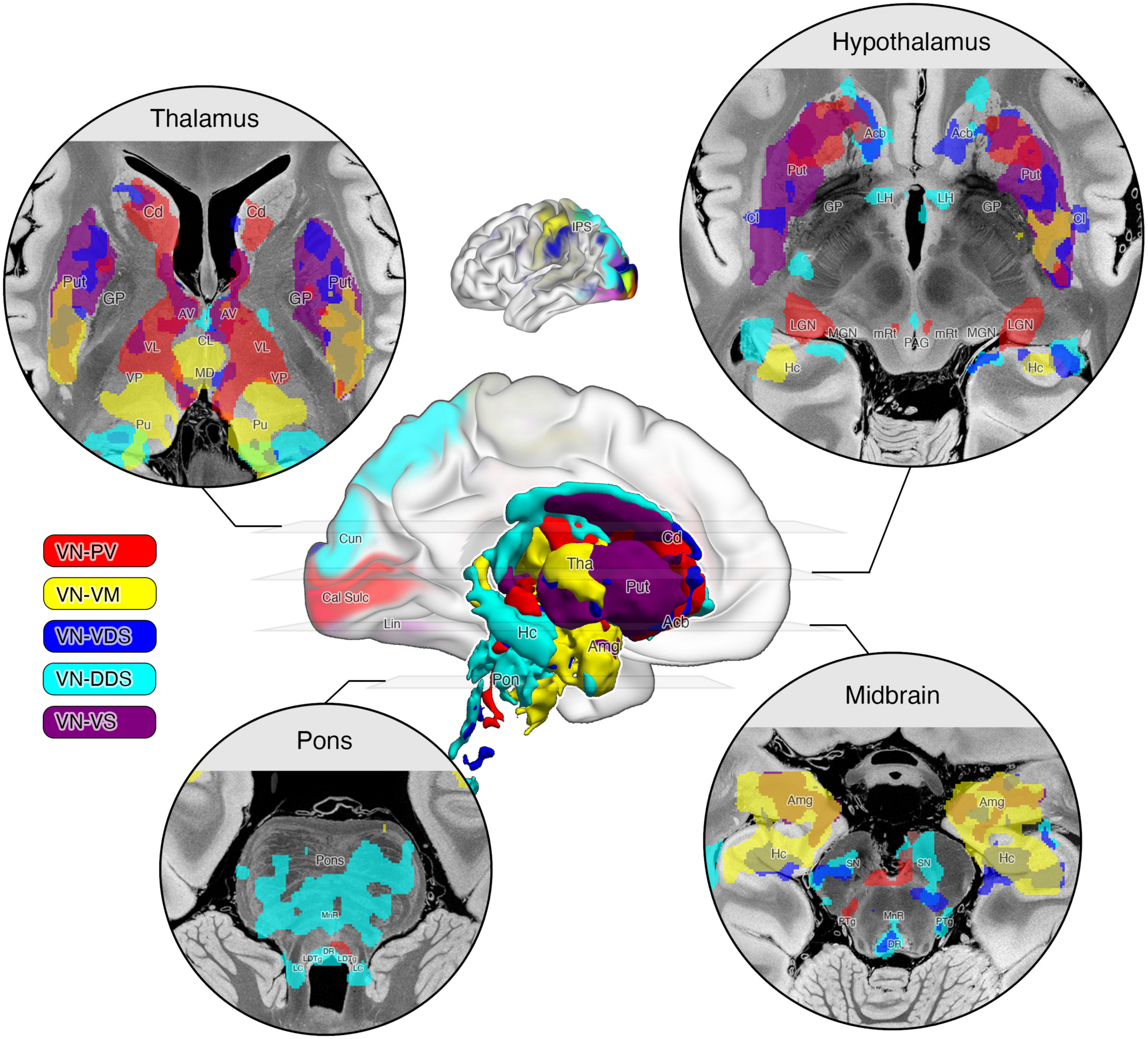
Visual network (VN) subnetworks. The primary visual (VN-PV) subnetwork is shown in red, the visual-motor (VN-VM) subnetwork is shown in yellow, the ventro-dorsal stream (VN-VDS) subnetwork is shown in blue, the dorso-dorsal stream (VN-DDS) subnetwork is shown in cyan, and the ventral stream (VN-VS) subnetwork is shown in purple. Refer to Fig. 1 for other notations.

#### The primary visual (VN-PV) subnetwork (red)

The primary visual subnetwork shows strong involvement in the primary visual cortex (V1) in the neighborhood of the calcarine sulcus. Key subcortical nodes of this subnetwork includes caudate, anterior ventral putamen, anterior (AV, VA), lateral (VLa, VLp, VPL), medial (CL, MDl, MDm), and dorsal (LD, LP) thalamic nuclei. In the posterior thalamic regions, LGN shows the strongest involvement across all networks in VN-PV. In addition, PAG is mildly involved among brainstem nuclei, and BNST is strongly involved among hypothalamic nuclei.

#### The visual-motor (VN-VM) subnetwork (yellow)

The VN-VM subnetwork involves the lateral part of V1, secondary (V2) and tertiary (V3) visual cortex, as well as primary sensorimotor area (M1/S1).

Strong involvement is observed mainly in the posterior section of the subcortical structures, such as hippocampus, amygdala, posterior putamen, posterior (PuA, PuL, PuM, VPL) and medial (CL, CM, MDm) thalamic nuclei. Mild involvement is also observed in the dorsal posterior caudate (tail). Neither the brainstem nor the hypothalamic nuclei are involved in this subnetwork, except for the pontine nuclei.

#### The ventro-dorsal stream (VN-VDS) subnetwork (blue)

This is the ventral branch of the dorsal stream. Cortical key nodes include V2, V3, SMG and AG along the IPS.

Subcortically, the caudate, accumbens, putamen, are hippocampus are strongly involved in VN-VDS. Key thalamic nodes include anterior (AV, VA), medial (CL, MDl, MDm) and dorsal (LD, LP) regions. In the brainstem, strong involvement is observed in DR and mild involvement is observed in PAG, LDTg, and PTg. BSNT is the only key node in the hypothalamic region for this subnetwork.

#### The dorso-dorsal stream (VN-DDS) subnetwork (cyan)

This is the dorsal branch of the dorsal stream. Cortically, VN-DDS spans V2, V3, V7 all the way along the SPG. Subcortically, VN-DDS involves dorsal caudate, accumbens, hippocampus, anterior (AV, VA), dorsal (LD, LP), medial (CL, MDm) and pulvinar (PuA, PuM, PuI, PuL) of the thalamus. Similar to VN-VDS, DR, PAG, LDTg, and PTg brainstem nuclei are involved in VN-DDS. And different from VN-VDS, the pons is strongly involved in this subnetwork. In addition, multiple hypothalamic nuclei are involved such as AHA, BNST, LH, MPO, NBM, RN, and SN.

#### The ventral stream (VN-VS) subnetwork (purple)

The ventral stream partially involves V1, as well as V2, V3, and ITG cortically. Subcortically, VN-VS is strongly involved in the caudate body, the entire putamen, and the amygdala. Among thalamic nuclei, VN-VS involves the dorsal anterior (AV, VA) and ventral posterior (LGN, PuI) regions. Brainstem and hypothalamic nuclei are not involved in this subnetwork.

### The higher-order cognitive networks (HOC)

The overall HOC, as shown in Fig. 4, spans regions in higher-order functional areas in the frontal, parietal and temporal lobes. Subcortically, the HOC strongly involves the thalamus, caudate, accumbens, putamen, globus pallidus, hippocampus, and amygdala, and multiple nuclei in the brainstem and hypothalamus.

**Fig. 4.**
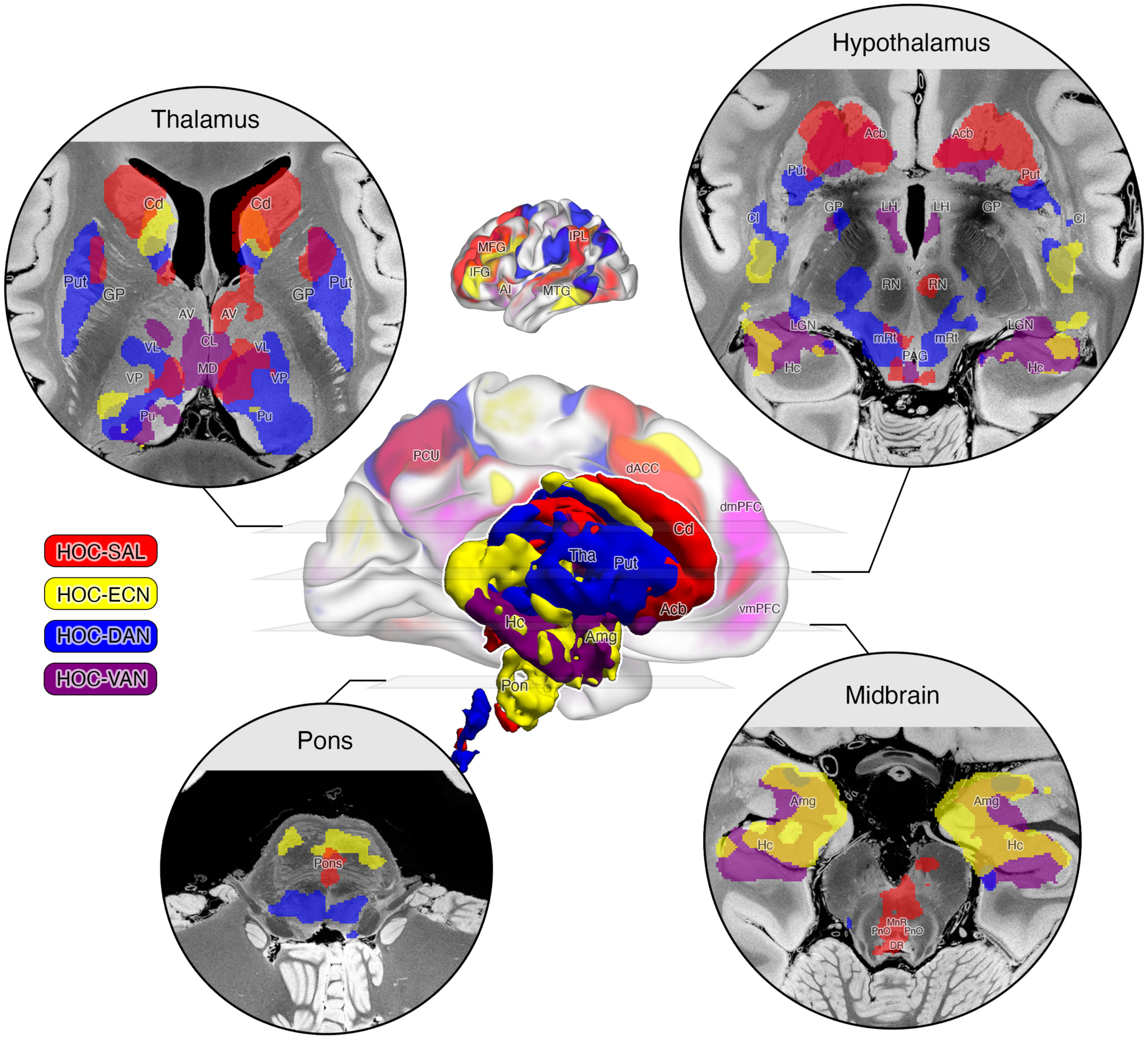
Higher-order cognitive (HOC) subnetworks. The salence (HOC-SAL) subnetwork is shown in red, the execuative control subnetwork (HOC-ECN) is shown in yellow, the dorsal attention subnetwork (HOC-DAN) is shown in blue, and the ventral attention subnetwork (HOC-VAN) is shown in purple. Refer to Fig. 1 for other notatolions.

#### The salence (HOC-SAL) subnetwork (red)

The cortical nodes of the HOC-SAL include dACC, anterior insula, pSMG, pSTG, and sfs. Subcortically, HOC-SAL mainly involves the dorsal and anterior caudate, accumbens, anterior putamen, and anterior globus pallidus. HOC-SAL also involves the dorsal sections of the thalamus (dorsal AV, VA, dorsal pulvinar, dorsal VLa, VLp, and LD). In the brainstem, HOC-SAL strongly involves midline nuclei such as DR, MnR, and midline pons. BNST is the most active node in the hypothalamus for this subnetwork.

#### The executive control (HOC-ECN) subnetwork (yellow)

The cortical nodes of the HOC-ECN include ifs, pits, AG laterally and two hot spots in SFG and PCC medially. Subcortically, HOC-ECN has strong involvement in the caudate inferior and posterior to the active caudate regions in HOC-SAL, hippocampus, and amygdala. HOC-ECN is also involved in the posterior tip of the putamen and the rostral posterior half of the thalamus from VPL, CM, MDl to LGN, MGN, L-SG, PuM, and PuI. Across the brainstem and hypothalamus, only the pons is highly involved.

#### The dorsal attention (HOC-DAN) subnetwork (blue)

The cortical key nodes of the HOC-DAN subnetwork include aSMG, pAG, and SPL along the ips and continue across the midline extended to PCu. It is also active in FEF and SEF in the posterior frontal lobe, pMTG in the posterior temporal lobe, as well as a small hotspot posterior to dACC. Subcortically, HOC-DAN is highly involved in the caudate right inferior and posterior to the active caudate regions in HOC-ECN, putamen and globus pallidus. In the thalamus, HOC-DAN involves the dorsal side of the posterior half including LP, VLp, VPL, CL, CM, MDm, MDl, and especially strong involvement in all pulvinar nuclei (PuA, PuM, PuL, PuI). HOC-DAN also involves PAG, mRt, PTg and pons in the brainstem and SN in the hypothalamus.

#### The ventral attention network (HOC-VAN) subnetwork (purple)

The cortical key nodes of the HOC-VAN subnetwork include IFG, anterior insula, pMTG, pITG, TPJ, pAG, amPFC, vPCC, pPCu, and sps. Subcortically, HOC-VAN is involved in the inferior caudate, accumbens, and ventral anterior putamen. It has strong involvement in the hippocampus and amygdala. In the thalamus, HOC-VAN involves medial and posterior nuclei such as CL, MdM, LGN, L-SG, PuM. Similar to HOC-DAN, HOC-VAN also involves PAG and mRt nuclei in the brainstem. In contrast to other HOC subnetworks, several hypothalamic nuclei are highly involved such as AHA, BNST, DM, LH, MM, SO, TM, and VMH.

#### Prediction of level of consciousness using individualized spatial maps of subnetworks

Approximately 25% of patients with severe brain injuries who appear unresponsive on the bedside behavioral examination are covertly conscious when assessed with task-based fMRI or EEG^41^. The presence of covert consciousness in the intensive care unit has been shown to predict the extent^42^ and pace^43^ of functional recovery. However, access to task-based technologies in clinical practice is limited to a small number of academic centers worldwide^44^. There is therefore great interest in identifying resting-state techniques, such as rs-fMRI, that are more widely available and have the potential to detect preserved brain networks essential for recovery of consciousness^45^.

As a proof-of-principle application of the functional atlases derived from the 1000 healthy controls, we mapped personalized subnetworks for each TBI patient in the RESPONSE dataset (n = 28)^46^. We showed that using a similarity measure between individualized subnetwork maps within the brainstem and the corresponding population-level “gold standard” functional atlases, along with age and sex as covariates, we were able to predict TBI patients’ consciousness levels, as measured by their total scores on the Coma Recovery Scale-Revised (CRS-R). Fig. 5 presents a scatter plot comparing the CRS-R total scores for 28 patients, assessed immediately before or after the rs-fMRI scan, with the predicted CRS-R total scores derived from a leave-one-out cross-validation procedure. The Pearson correlation between these two measures was r=0.46 (uncorrected p=0.01), indicating a moderate, statistically significant association.

**Fig. 5.**
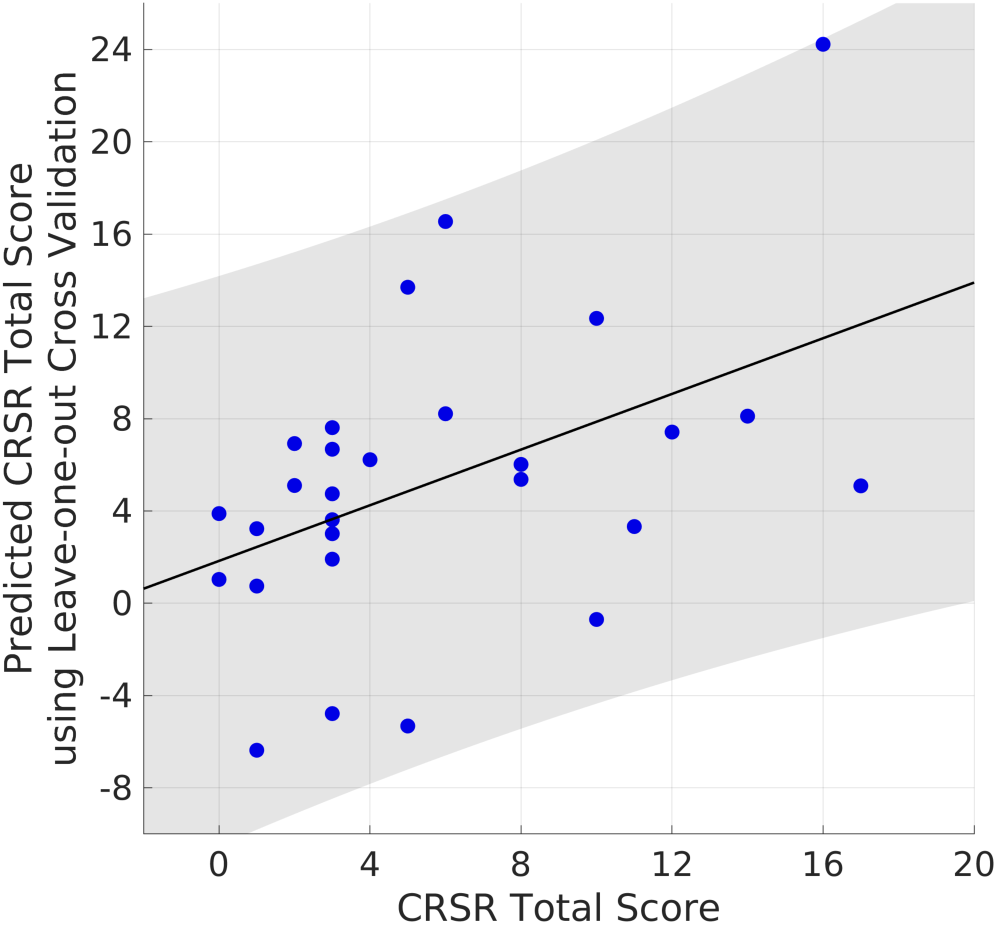
Scatter plot between the measured CRS-R total scores (x axis) and the predicted CRS-R total scores from 28 TBI patients in the RESPONSE cohort. The shaded area indicates 95% confidence interval for the best linear fit between the two.

When we extended this analysis to other brain regions, we found that the functional connectivity of the thalamus also predicted CRS-R total scores (r=0.44, p=0.02). In contrast, similarity measures derived from the entire subcortical region (r=0.08, p=0.69), the cortical region (r=0.03, p=0.88), and the whole brain (r=-0.06, p=0.75) did not predict CRS-R total scores.

Further, by fitting a single ordinary least squares regression model using all 28 patients, we identified the following predictor variables that significantly contributed to the prediction of CRS-R total scores: age (p=0.014), HOC-VAN (p=0.012), HOC-SAL (p=0.021), and SMN-Face (p=0.042). VN-VS (p=0.054) and DMN-MTL (p=0.087) show trends towards significance.

## Discussion

Subcortical functional mapping is crucial for advancing our understanding and treatment of a wide range of neurological and psychiatric disorders. In this study, we demonstrated that the subcortical connections of large-scale cerebral cortical networks are highly reproducible in the healthy human brain. Further, we employed a novel analytic technique – NASCAR – designed to overcome prior methodological barriers in separating overlapping networks, to enable fractionation of large-scale brain networks into subnetworks with distinct subcortical connectivity patterns. These subnetworks exhibited differential patterns of subcortical connectivity, yet they all demonstrated substantial spatial overlap, particularly within widely connected hub nodes of the caudate, putamen, hippocampus, and thalamus. Cortical counterparts, by contrast, showed varying degrees of overlap, with some subnetworks, such as the DMN and HOC subnetworks, demonstrating considerable overlap, while others, such as the SMN subnetworks, were more spatially segregated. Collectively, these observations indicate a highly conserved and spatially overlapped subcortical functional architecture in the human brain, providing opportunities for elucidating the pathogenesis of neuropsychiatric diseases and for developing new treatments aimed at modulating subcortical brain networks.

The fractionation of large-scale human brain networks in this study was inspired by prior efforts to fractionate the cortical components of these networks, in particular the DMN^4^. However, in contrast to the conventional seed-based method, we approached this problem through the lens of whole-brain network analyses, which allowed us to extend fractionation into the human subcortex, while also providing insights into fractionation of overlapped cortical networks. Whereas prior DMN connectivity studies suggested two subnetworks – the DMN-MTL and the DMN-dmPFC (for which we use nomenclature proposed by Andrews-Hanna and colleagues^4^) – we identified a third DMN component, for which we propose the name DMN-vPCC, based on the strong connectivity to the ventral posterior cingulate cortex. The DMN-vPCC appears to be the primary DMN subnetwork that connects with key arousal centers in the rostral brainstem, including the noradrenergic LC and glutamatergic mRT. While the functional relevance of DMN-vPCC, in comparison to DMN-MTL and DMN-dmPFC, awaits further investigation, the subcortical involvement of DMN is consistent with prior findings,^7,47,48^ and our results provide a spatially contiguous view of each subnetwork at unprecedented neuroanatomic resolution.

For the SMN, VN, and HOC, although some observations were aligned with results from structural and/or ex vivo studies (e.g., LGN is a subcortical relay node for the primary visual network^49,50^), there is limited prior evidence to which functional subcortical fractionation findings can be compared. In this light, the findings here can be considered hypothesis-generating.

As a proof-of-principle, we showed that the higher similarity of individual subnetworks within the brainstem (and thalamus) from TBI patients to the gold standard established from the healthy controls, the higher their consciousness levels. This predictability does not hold for overall subcortical regions, cortical regions, or the whole brain, suggesting that the subcortical subnetworks identified here have specific relevance to recovery of consciousness after severe brain injury. Interestingly, multiple subcortical subnetworks beyond the DMN-MTL, including HOC-VAN, HOC-SAL, SMN-Face, and VN-VS, contributed to predicting CRS-R total scores, consistent with prior cortical connectivity studies showing that the DMN may be necessary, but not sufficient, for generating human consciousness^51^.

A limitation of this work, common across all rsfMRI studies, is the lack of a ground truth for large-scale brain networks. This limitation not only contributes to ambiguities in nomenclature^23^ but also presents a significant challenge for validation. We tackled this problem using a bootstrapping strategy via subject resampling with replacement and repeated NASCAR analyses. Our results showed that the identified large-scale brain networks are highly reproducible. Additionally, we note that the spatial topology of these networks is naturally inherited from the data, rather than being a technical artifact of the analytical method. For instance, the three DMN subnetworks span the caudate nucleus in a spatially neighbored yet well-organized manner as shown in the 3D illustratation in Fig. 1, although there is no mathematical constraint to force them to be organized in such a way.

Although we observed substantial overlap in the identified brain networks/subnetworks, particularly in the subcortical regions, we acknowledge the inherent limitations associated with linking mesoscale rsfMRI data to microscale neuroanatomic and electrophysiologic data^52,53^. For this reason, we caution that the interpretation of spatial overlap in this study be considered as a mesoscale property of the brain, pending future exploration and validation in correlative experiments performed at microscale^54^. The limited spatial resolution and SNR of the fMRI data, as well as the imperfect inter-subject registration, may constrain our ability to accurately delineate specific functional areas and their boundaries. Indeed, networks, at both cortical and subcortical level, have been shown to be organized into multiple functionally specialized regions in a hierarchical manner within individuals using intensive and repeated fMRI acquisition^55–57^.

The current study leveraged 1,000 scans from the HCP and investigated the subcortical functional connectivity of large-scale brain networks that are common across the healthy population. It is important to note that this study did not provide a comprehensive list of resting-state networks, nor did it investigate task-based networks (e.g., the auditory and language network^55^ was not identified). Additionally, individuals with different age ranges or brain pathologies may exhibit distinct brain network configurations compared to the young adult cohort from the HCP. Thus, our findings will require further test-retest validation and external validation using other datasets.

An additional limitation of the current approach is the heavy computational burden associated with the NASCAR method, which may require significant time and robust computational resources to replicate the experiments with similarly sized or larger datasets. Moroever, the results presented here were derived using research-grade fMRI data, and further investigation is needed to assess their applicability to clinical scans. From the standpoint of clinical translation, the present study did not explore personalized brain network mapping, an approach that holds considerable potential for precision medicine and represents a promising direction for future research.

Despite these limitations, this work advances knowledge about the functional network architecture of the human brain and extends our understanding of spatially overlapped brain networks from the cerebral cortex to the subcortex. We fractionated and identified 15 spatially overlapped and temporally correlated subnetworks that can be categorized into 4 large-scale brain networks and are highly reproducible across healthy human brains. We revealed that the subcortical functional connectivity patterns of these subnetworks exhibited distinct yet overlapped spatial configurations and were intrinsically organized in neuroanatomic relationships that could be derived from the fMRI data, rather than being attributable to the mathematical model. Further, we demonstrated, as a proof-of-principle, that the similarity between individual subnetwork maps and the gold standard functional atlases within the brainstem predicts the consciousness levels in TBI patients. The subcortical network maps generated here, which we release to the academic community, thus create opportunities to elucidate pathophysiologic mechanisms and develop new neuromodulatory treatments for a broad range of neuropsychiatric disorders.

## Materials and Methods

### Datasets

#### The Human Connectome Project (HCP) dataset

The minimally preprocessed 3 Tesla rs-fMRI data of 1000 subjects from the Human Connectome Project (HCP)^58,59^ were used in this study. Each subject has two sessions of rs-fMRI scans with two different phase encoding directions (LR, RL). We used data from the LR direction only to minimize the inter-subject registration error due to the potential residual differences in EPI distortion after correction^60^. Only the first session of data was used for computational feasibility. Hence, 1000 rs-fMRI scans, one from each subject, were used in this project in total. The acquisition parameters include TR=720ms, TE=33ms, and 2 mm isotropic spatial resolution. The rs-fMRI data were co-registered to the HCP MNI atlas through the T1-weighted (T1w) image using a single concatenated deformation. Each rs-fMRI session lasted 15 min with 1200 frames in total. The volumetric data were resampled onto the native cortical surfaces extracted from each subject’s T1w image, and then co-registered to the HCP fsLR surfaces^58^. All analyese were performed in this HCP grayordinate space. We did not apply any additional spatial smoothing beyond the 2 mm full width half maximum isotropic Gaussian smoothing that was used in the minimal preprocessing pipeline^58^, to avoid blurring across functional boundaries^61–63^.

#### The RESPONSE dataset

We used a clinical dataset acquired in the “REsting and Stimulus-based Paradigms to detect Organized NetworkS and predict Emergence of consciousness” (RESPONSE) study^45^. 61 acute TBI patients admitted to the Neurosciences ICU, Multidisciplinary ICU, and Surgical ICU at Massachusetts General Hospital (MGH) between September 2018 and December 2022 were enrolled in the study. Each patient was assessed with the Coma Recovery Scale-Revised (CRS-R)^64^ – a standardized behavioral measure that is recommended by multiple professional organizations for assessment of DoC^65,66^ – immediately prior to the imaging session. MRI data were acquired with a 32-channel head coil on a 3 Tesla Skyra MRI scanner (Siemens Healthineers; Erlangen, Germany) located in the MGH Neurosciences ICU. A 3D T1-weighted multi-echo magnetization prepared gradient echo (MEMPRAGE) anatomical images were acquired at 1mm isotropic resolution. The rs-fMRI data were acquired using a BOLD sequence with TE=30ms, TR=1250ms, a simultaneous multislice (SMS) factor of 4, and a 2 mm isotropic resolution. 33 out of 61 patients successfully completed both the CRS-R assessment and MR data acquisition in their acute phase^46^. We used FreeSurfer v7.4.0^67^ for anatomical image processing and surface generation. We used fMRIPrep^68^ to preprocess the BOLD data and resampled them onto the grayordinate space which is compatible with the HCP dataset. 5 out of the 33 patients did not complete the FreeSurfer or the fMRIPrep pipeline due to either insufficient quality of the MRI data or brain abnormality (e.g. presence of large lesions), resulting in 28 patients in the final cohort used in this work.

### Fractionation and network identification using NASCAR

#### The NASCAR method

We applied the Nadam-Accelerated SCAlable and Robust (NASCAR) canonical polyadic decomposition algorithm^7,38,39^ to separate large-scale brain networks from the group rs-fMRI data. Fig. S1 (a) illustrates this concept. Two choirs are singing different songs simultaneously in a room. The individuals in the red choir are performing one song, while those in the yellow choir are singing a different one. Two microphones in the room capture the mixed voices. The choir members are spatially dispersed and intermingled within the room. In analogy to this scenario, within the context of brain network separation/identification, the red and yellow choirs represent two distinct brain subnetworks, each exhibiting its own neuronal activity (the songs). The mixed recordings correspond to the acquired fMRI data. NASCAR is designed to recover the spatially overlapping (intermingled individuals) and temporally correlated (simultaneous singing) brain activities associated with each subnetwork (true signals) from the mixed recordings (acquired fMRI data).

Specifically, we first applied the group BrainSync algorithm^69,70^ to align the time series from each subject to a common temporal space. To increase the signal contribution from subcortical regions (the low-rank NASCAR optimization is biased towards regions that have more voxels/vertices) and enhance computational feasibility, we excluded the data from the cerebellum and downsampled the data from cortical regions by a factor of three^38^. Then the synchronized rs-fMRI data were concatenated along the third (subject) dimension to form a data tensor *X* ∈ ℝ*^V^*^×*T*×*S*^, where *V* ≈ 36*K* is the number of vertices (space), *T* = 1,200 is the number of frames (time), and *S* = 1,000 is the number of subjects (subject), Fig. S1 (b). Similar to independent component analysis (ICA) or principal component analysis (PCA) in 2D cases, NASCAR finds low-rank components from 3D tensors. However, unlike ICA or PCA where the components are required to be either independent or orthogonal to each other, NASCAR does not impose any constraints, which is more physiologically plausible for brain networks^20,71^. We ran NASCAR on the data tensor with a rank up to 30 as fMRI data were low-rank in nature^31,33,72^. Instead of “networks” or “subnetworks”, we specifically use the term “components” to describe the NASCAR results because not all low-rank components are aligned with well-known large-scale brain networks. The recognition of networks and their subnetworks is described below after the bootstrapping.

#### Bootstrapping

The NASCAR method has superior robustness compared to traditional tensor decomposition algorithms due to its warm initialization strategy^73^. In this work, we further improved the robustness of brain network identification using bootstrapping. Specifically, we ran NASCAR on the 1000 subjects’ rs-fMRI data 100 times. Each time we obtained 30 low-rank components with a random resampling over the subject with replacement. We gathered all 3000 components from the 100 runs. We used the cross-correlation *C* ∈ ℝ^3000×3000^ of the spatial maps as the similarity measures between components. We applied the uniform manifold approximation and projection (UMAP)^74^ together with the hierarchical density-based spatial clustering of applications with noise (HDBSCAN) algorithm to group the components into 30 clusters because UMAP better preserves topological relationships. We also applied the curvilinear component analysis (CCA)^75^ to embed the spatial maps from the ≈ 96*K* dimensional space to a 2-dimensional (2D) space for visualization of their relative relationships because CCA preserves the distances between components (the closer the spatial maps are in the higher dimensional space, the closer they are in the embedded 2D space and vice versa). Fig. S2 shows the CCA-embedded spatial maps color-coded using the UMAP+HDBSCAN clustering labels. Each dot represents a spatial map from the 3000 bootstrapped NASCAR results. A tighter cluster indicates a higher robustness during the NASCAR decomposition (i.e., if a certain component is obtained from the analysis, it had almost the same solution every time the analysis was run). Fig. S3 shows the CCA-embedded spatial maps identical to Fig. S2 except for the color coding. The colors in this figure represent the reproducibility scores, which are measured by the count of the presence of a component across all 100 bootstrap runs (a score of 100 indicates this component is found in the result from every single 100 runs, hence highly reproducible). Figs S2 and S3 collectively show that the primary brain networks, such as VN and SMN, are highly robust and reproducible across the population and the higher-order cognitive networks, such as DMN, SAL, ECN, DAN, and VAN, have moderate-to-high robustness and reproducibility.

#### Network/Subnetwork identification and recognition

We identified 15 brain networks via posterior visualization of the components and their relationships as with any other blind-source separation methods (e.g., ICA, PCA). Specifically, for each cluster from the boostrapping results, we took the mean of all points to have a single spatial map for that cluster. We then separated the cortical sections of the mean spatial map from the result and plotted them on the left and right cortical surfaces, respectively. We also looked into the inter-cluster relationships by computing and visualizing the correlation matrices in space, time, subjects, and spectra, as shown in Fig. S4. The 15 subnetworks were recognized and grouped into 4 large-scale network categories iteratively using the combination of 1) how well the spatial map aligns with neuroanatomy priors (e.g., SMN-Hand shows activation in the superior segment of pre- and post-central regions); 2) how well the spatial map aligns with prior literatures (e.g., DMN subnetworks should have activations in PCC, mPFC, etc.); and 3) the relationship among subnetworks through the correlation matrices (e.g., VN subnetworks have high spatial correlations with each other as they all occupy the occipital lobe, the HOC subnetworks have high positive correlations with each other but also high anti-correlation with the DMN subnetworks temporally, etc.)

#### Separation of subcortical component of the networks

We highlight the key difference in how brain networks are identified in the subcortex. In contrast to commonly used seeded correlation analysis where seeds are placed in two ROIs separately (e.g., one in a subcortical region and another one in a cortical region), NASCAR finds the subcortical maps simultaneously with the cortical maps as a whole-brain network analysis. Fig. S1 illustrates this procedure. Each network/subnetwork that NASCAR identified is an outer product of a spatial map (vertical bar), a time series (horizontal), and a subject participation mode (oblique). The spatial maps of these low-rank components lie in the same CIFTI space as with the rs-fMRI data. In Fig. S1, the three DMN subnetworks are used for illustrative purposes. The spatial maps, in the zoomed view on the right-hand side of Fig. S1, include both cortical and subcortical locations. The top one-third of the vector represents the spatial map on the left hemisphere that can be mapped back to the left cortical surface for visualization. Similarly, the middle-one third is for the right hemisphere. The bottom one-third of the vector is the spatial map on the subcortical structures and is mapped back to the MNI space in a volumetric representation. NASCAR finds the spatial maps for the whole brain (the entire vector) at the same time. By visualizing the cortical spatial maps (top two-third), we recognized the subnetworks as described above. Thus, the last one-third of each spatial map must correspond to its corresponding subnetworks, respectively. The main results reported in this work are the separation and visualization of the subcortical parts of the spatial maps in the volumetric space.

#### Visualization

For each identified subnetwork, we thresholded at the top 5% cut-off to form a binary map and color-coded as shown in Figs. 1-4. For each large-scale brain network group (DMN, SMN, VN, HOC), we plotted the cortical maps of its subnetworks on a slightly smoothed pial surface (in the common fsLR space) of the exemplar subject (100307) provided by the HCP as shown in the center panel (semi-transparent lateral view of the left cortical surface) and top miniaturized panel (medial view of the left cortical surface) of Figs. 1-4. We overlaid the subcortical maps of its subnetworks in the volumetric MNI space on a 7T 100um ex vivo MRI scan^76^ for precise anatomical analyses as shown in the four corner panels of Figs. 1-4. We note that, due to the large number of subnetworks in total, we reused the colors (e.g. red, yellow, blue) in each figure to represent their subnetworks, i.e., for example, the red subnetwork in Fig. 1 represents DMN-MTL, which is different and independent from the red subnetwork SMN-Face shown in Fig. 2.

For the purpose of visualization in a surface view of the subcortical subnetworks only, we additionally smoothed the binary maps volumetrically using a 3D Gaussian filter with an isotropic window size of 5 voxels and a kernel standard deviation of 2. This Gaussian smoothing discarded isolated voxels or small clusters in the next tessellation step. We then tessellated the smoothed subcortical map for each subnetwork (‘mri_tessellate’) and performed an additional surface smoothing (‘mris_smooth’), both using FreeSurfer^67^. The resulting surfaces of the subnetworks were rendered in Freeview (the viewer for FreeSurfer) and color-coded using the same schema as with the volumetric subcortical maps. A single screenshot was taken for each network from the medial view and embedded, together with the medial view of the left cortical surface, in the center panel of Figs. 1-4. An interactive visualization of these surface views of the subcortical subnetworks was created and rendered on this website for easy access: https://subcorticalfractionation.netlify.app.

### Clinical application

To illustrate a potential clinical application of the subcortical functional atlases derived from 1000 healthy controls from HCP, we mapped personalized subnetworks for each TBI patient in the RESPONSE dataset (n = 28) using their individual rsfMRI scans^46^. For each subnetwork, we extracted the spatial map within the brainstem region (defined by the FreeSurfer aseg label brainstem + ventral diencephalon) and assessed its similarity to the corresponding subnetwork in the atlases. This produced a scalar measure for each subnetwork in each patient (e.g., for the DMN-MTL, this value indicates the degree of similarity between the patient’s individual DMN-MTL map and the population-level “gold standard” DMN-MTL map derived from 1000 HCP healthy controls). To control for potential confounding effects, we incorporated age and sex as covariates, appending them to the similarity measures for each patient. Thus, the independent variables consisted of 28 patients, each characterized by 17 predictors (15 network similarity measures, age, and sex). The CRS-R score for each patient served as the dependent variable.

Experiment 1: We evaluated the predictive power and generalizability of the subcortical network-based measures using a leave-one-out cross-validation approach. Specifically, for each of the 28 patients, we excluded one patient, trained an ordinary least squares regression model on the remaining 27 patients, and predicted the CRS-R score for the excluded patient. This process was repeated for all patients, and the predicted CRS-R scores were compared to the observed CRS-R scores. The relationship between the observed and predicted CRS-R scores was quantified using Pearson correlation.

Experiment 2: We extended the investigation to determine whether network similarity in regions beyond the brainstem could also predict CRS-R scores. In this experiment, we repeated the procedure from Experiment 1, but evaluated the predictive power of individualized subnetwork spatial maps derived from four distinct brain regions: 1) the whole brain, 2) the cortical regions only, 3) the entire subcortical region, and 4) the thalamus.

Experiment 3: To identify which subnetworks or confounding factors most strongly contribute to predicting CRS-R scores, we fitted a single ordinary regression model using all 28 patients and assessed the statistical significance of each predictor variable.

## Acknowledgment

Support for this research was provided in part by the BRAIN Initiative Cell Atlas Network (BICAN; U01MH117023, UM1MH134812 and UM1MH130981), the Brain Initiative Brain Connects consortium (U01NS132181, UM1NS132358), National Institutes of Health Director’s Office (DP2HD101400), National Institute for Neurological Disorders and Stroke (R21NS109627, R01NS138257, R01NS128961, RF1NS115268, U24NS135561, R01NS070963, R01NS083534, R01NS105820, R25NS125599, R01NS127892, UM1NS132358), National Institute for Biomedical Imaging and Bioengineering (R01EB023281, R21EB018907, R01EB019956, P41EB030006), National Institute on Aging (R21AG082082, R01AG064027, R01AG016495, R01AG070988), National Institute of Mental Health (UM1MH130981, R01MH123195, R01MH121885, RF1MH123195, R01MH130666, R01MH113929), and Department of Defense (W81XWH2210999). Additional support was provided by the NIH Blueprint for Neuroscience Research (U01MH093765), part of the multi-institutional Human Connectome Project, the Chen Institute MGH Research Scholar Award, the MIT/MGH Brain Arousal State Control Innovation Center (BASCIC) project, the Schilling Foundation, and the German Research Foundation (Deutsche Forschungsgemeinschaft, CRC-1451, 431549029). Much of the computation resources required for this research was performed on computational hardware generously provided by the Massachusetts Life Sciences Center (https://www.masslifesciences.com/). This work was also made possible by the resources provided by Shared Instrumentation Grants S10RR023401, S10RR019307, and S10RR023043.

## Competing interests

B. F. is an advisor to DeepHealth, a company whose medical pursuits focus on medical imaging and measurement technologies. BF’s interests were reviewed and are managed by Massachusetts General Hospital and Mass General Brigham in accordance with their conflict of interest policies.

A. H. reports lecture fees for Boston Scientific, is a consultant for Modulight.bio, was a consultant for FxNeuromodulation and Abbott in recent years and serves as a co-inventor on a patent granted to Charité University Medicine Berlin that covers multisymptom DBS fiberfiltering and an automated DBS parameter suggestion algorithm unrelated to this work (patent #LU103178).

## Data and code availability

The data used in this study are publicly available from the Wash U/U Minn component of the Human Connectome Project, Young Adult Study at https://www.humanconnectome.org/study/hcp-young-adult. For research purposes, we will release the functional atlases and the code publicly once the manuscript is accepted.

## Supplementary Materials

**Table S1.** Subnetwork activity level for overall subcortical structures. An upper case “X” indicates there is a strong activity, a lower case “x” indicates there is a moderate/mild activity, and a blank cell indicates there is minimal/no activity. Blue shapes are used to separate the four groups of networks for easy visualization.

|  | Caudate | Accumbens | Putamen | Globus Pallidus | Hippocampus | Amygdala | Clastrum |
| --- | --- | --- | --- | --- | --- | --- | --- |
| DMN-MTL | X | x |  |  | X | X |  |
| DMN-dmPFC | X |  |  |  | x |  |  |
| DMN-vPCC | x | X | x |  | X |  |  |
| SMN-Face | x |  | X | X |  |  | x |
| SMN-Hand | x |  | x |  | X |  |  |
| SMN-Foot | x |  | X |  |  |  | x |
| VN-PV | X |  | x |  |  |  |  |
| VN-VM | x |  | X |  | X | X | x |
| VN-VDS | X | X | X |  | X |  | x |
| VN-DDS | X | X |  |  | X |  |  |
| VN-VS | X |  | X |  |  | X | x |
| HOC-SAL | X | X | x | x |  |  |  |
| HOC-FPN | X |  | x |  | X | X | x |
| HOC-DAN | x |  | X | x |  |  | x |
| HOC-VAN | x | X | x |  | X | X |  |

**Table S2.**
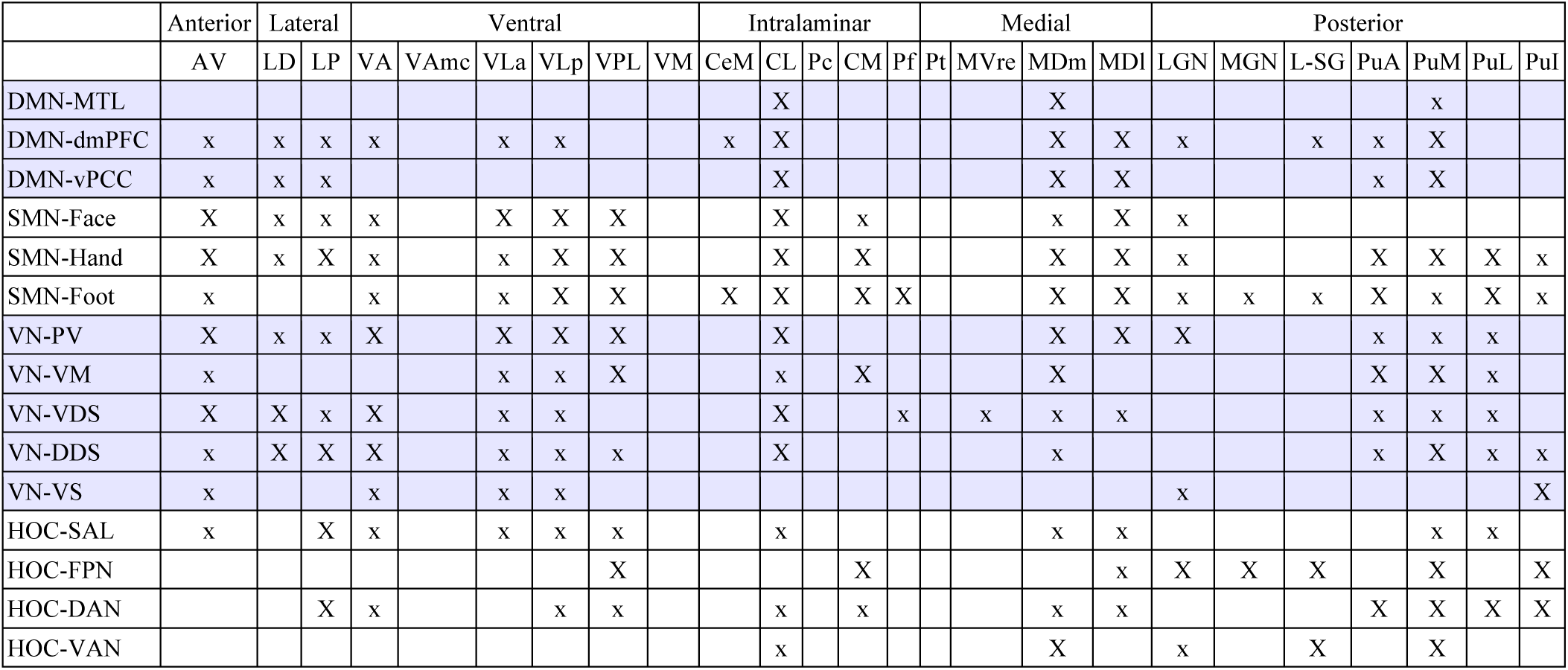
Subnetwork activity level for thalamic nuclei. See Table 1 for the notations.

**Table S3.** Subnetwork activity level for brainstem nuclei. See Table 1 for the notations.

|  | DR | MnR | PAG | VTA | LC | LDTg | mRt | PBC | PnO | PTg | Pons |
| --- | --- | --- | --- | --- | --- | --- | --- | --- | --- | --- | --- |
| DMN-MTL | x | X |  | x |  | X |  |  | X |  |  |
| DMN-dmPFC |  |  |  | x |  |  |  |  |  | x | X |
| DMN-vPCC | X | X |  | x | X | X | x | X | X |  |  |
| SMN-Face |  |  |  |  | X |  |  | X |  |  | x |
| SMN-Hand |  |  |  |  |  |  |  |  |  |  | x |
| SMN-Foot |  |  | x |  |  |  | x |  |  |  | x |
| VN-PV |  |  | x |  |  |  |  |  |  |  |  |
| VN-VM |  |  |  |  |  |  |  |  |  |  | x |
| VN-VDS | X |  | x |  |  | x |  |  |  | x |  |
| VN-DDS | X |  | x |  |  | X |  |  |  | x | X |
| VN-VS |  |  |  |  |  |  |  |  |  |  |  |
| HOC-SAL | X | x |  |  |  |  |  |  |  |  | x |
| HOC-FPN |  |  |  |  |  |  |  |  |  |  | x |
| HOC-DAN |  |  | X |  |  |  | X |  |  | x | x |
| HOC-VAN |  |  | x |  |  |  | x |  |  |  |  |

**Table S4.** Subnetwork activity level for hypothalamic nuclei. See Table 1 for the notations.

|  | AHA | AN | BNST | DP | DM | LH | MM | MPO | NBM | Pa | Pe | PH | RN | SC | SN | SO | STN | TM | VMH | ZI |
| --- | --- | --- | --- | --- | --- | --- | --- | --- | --- | --- | --- | --- | --- | --- | --- | --- | --- | --- | --- | --- |
| DMN-MTL | x | x | x |  |  | x | x |  |  |  |  |  |  |  |  | X |  | X | x |  |
| DMN-dmPFC |  |  |  |  |  |  |  |  |  |  |  |  |  |  | X |  |  |  |  |  |
| DMN-vPCC |  |  |  |  |  | x | X |  |  |  |  |  |  |  | x |  |  |  |  |  |
| SMN-Face |  |  |  |  |  |  |  |  | x |  |  |  |  |  |  |  |  |  |  |  |
| SMN-Hand |  |  |  |  |  |  |  |  |  |  |  |  |  |  |  |  |  |  |  |  |
| SMN-Foot |  |  |  |  |  |  |  |  |  |  |  |  |  |  |  |  |  |  |  |  |
| VN-PV |  |  | X |  |  |  |  |  |  |  |  |  |  |  |  |  |  |  |  |  |
| VN-VM |  |  |  |  |  |  |  |  |  |  |  |  |  |  |  |  |  |  |  |  |
| VN-VDS |  |  | X |  |  |  |  |  | x |  |  |  |  |  | x |  |  |  |  |  |
| VN-DDS | x |  | x |  |  | X |  | x | x |  |  |  | x |  | x |  |  |  |  |  |
| VN-VS |  |  |  |  |  |  |  |  |  |  |  |  |  |  |  |  |  |  |  |  |
| HOC-SAL |  |  | X |  |  |  |  |  |  |  |  |  |  |  |  |  |  |  |  |  |
| HOC-FPN |  |  |  |  |  |  |  |  |  |  |  |  |  |  |  |  |  |  |  |  |
| HOC-DAN |  |  |  |  |  |  |  |  |  |  |  |  |  |  | X |  |  |  |  |  |
| HOC-VAN | X |  | x |  | x | X | X |  |  |  |  |  |  |  |  | X |  | x | x |  |

**Fig. S1.**
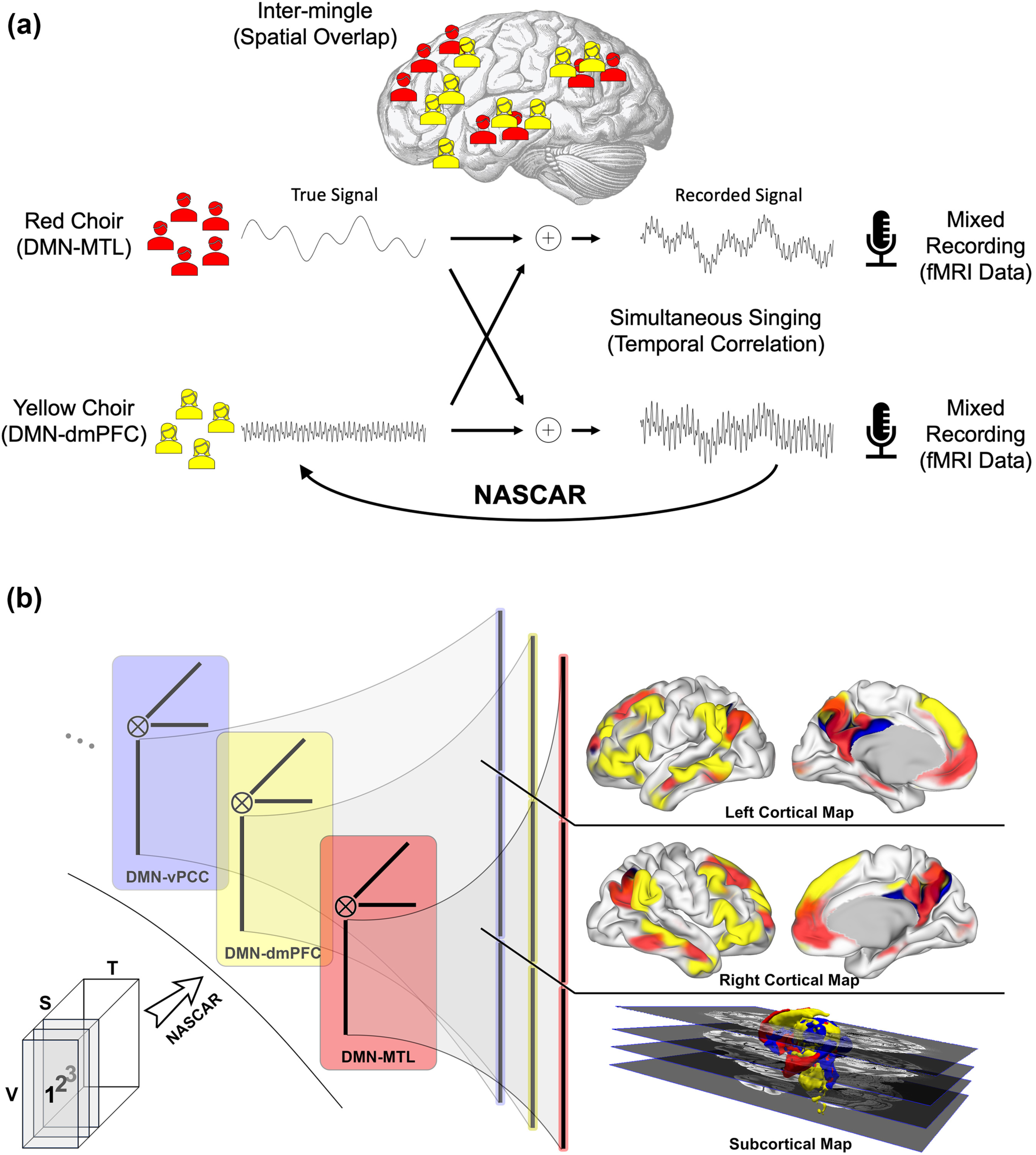
The NASCAR method for identification of overlapping subnetworks of the brain. (a) Illustration of the spatially overlapped and temporally correlated networks that NASCAR is designed to separate using a choir analogy. Two choirs are singing different songs simultaneously in a room. The individuals in the red choir are performing one song, while those in the yellow choir are singing a different one. Two microphones in the room capture the mixed voices. The choir members are spatially dispersed and intermingled within the room. In analogy to this scenario, within the context of brain network separation/identification, the red and yellow choirs represent two distinct brain subnetworks, each exhibiting its own neuronal activity (the songs). The mixed recordings correspond to the acquired fMRI data. NASCAR is designed to recover the spatially overlapping (intermingled individuals) and temporally correlated (simultaneous singing) brain activities associated with each subnetwork (true signals) from the mixed recordings (acquired fMRI data). (b) Data formulation, NASAR decomposition, whole-brain network identification, and CIFTI representation. Resting-state fMRI data are arranged in a 3D tensor with V number of vertices in the CIFTI spatial representation (space), T number of frames (time), and S number of subjects. NASCAR finds low-rank components, each of which is an outer product of a spatial map (vertical bar), a time series (horizontal), and a subject participation mode (oblique). Three DMN subnetworks are used for illustration purpose. The spatial maps identified by the NASCAR method include both cortical and subcortical locations as illustrated on the right hand side. The zoom-in spatial maps shows the arrangement of spatial data in the CIFTI format: the top one third of the vecter represents the spatial map on the left hemisphere that can be mapped back to the left cortical surface for visualization. Similarly for the right hemisphere in the middle one third. The bottom one third of the vector is the spatial map on the subcortical structures and is mapped back to the MNI space in a volumetric representation.

**Fig. S2.**
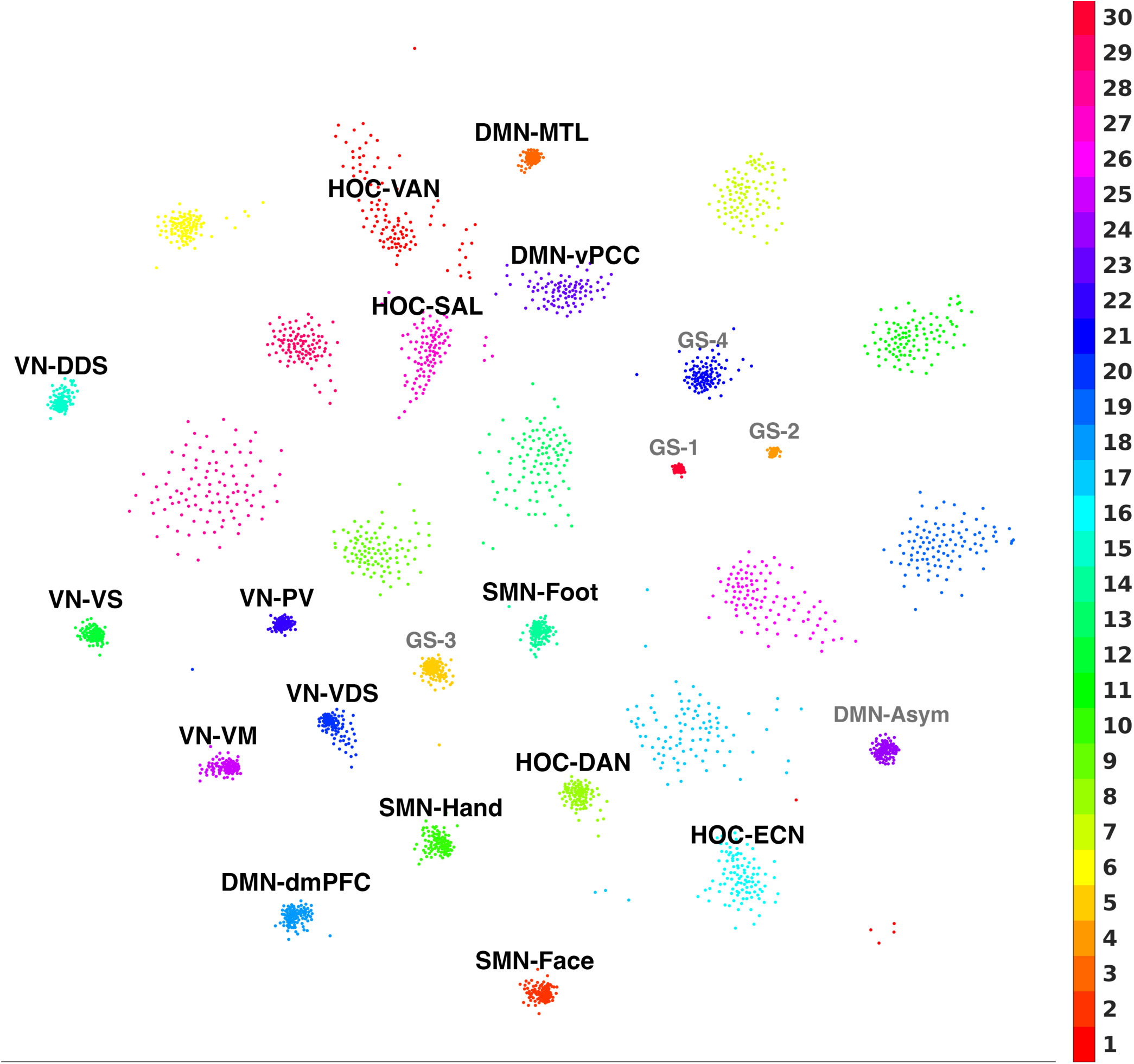
CCA-embedded spatial maps with UMAP+HDBSCAN clusters. Each dot represents a spatial map from the 3000 bootstrapped NASCAR results. The color bar indicate the discrete colors, one for each UMAP+HDBSCAN clusters. The 15 recognized subnetworks are labeled in bold black. Consistent with our previous work (Li et al., 2023), four global signals (GS) and one additional asymmetric DMN component (DMN-Asym) were also found highly reproducible across subjects and boostrap runs. They are not included in the subcortical analyses for this work but labeled in light gray for completeness.

**Fig. S3.**
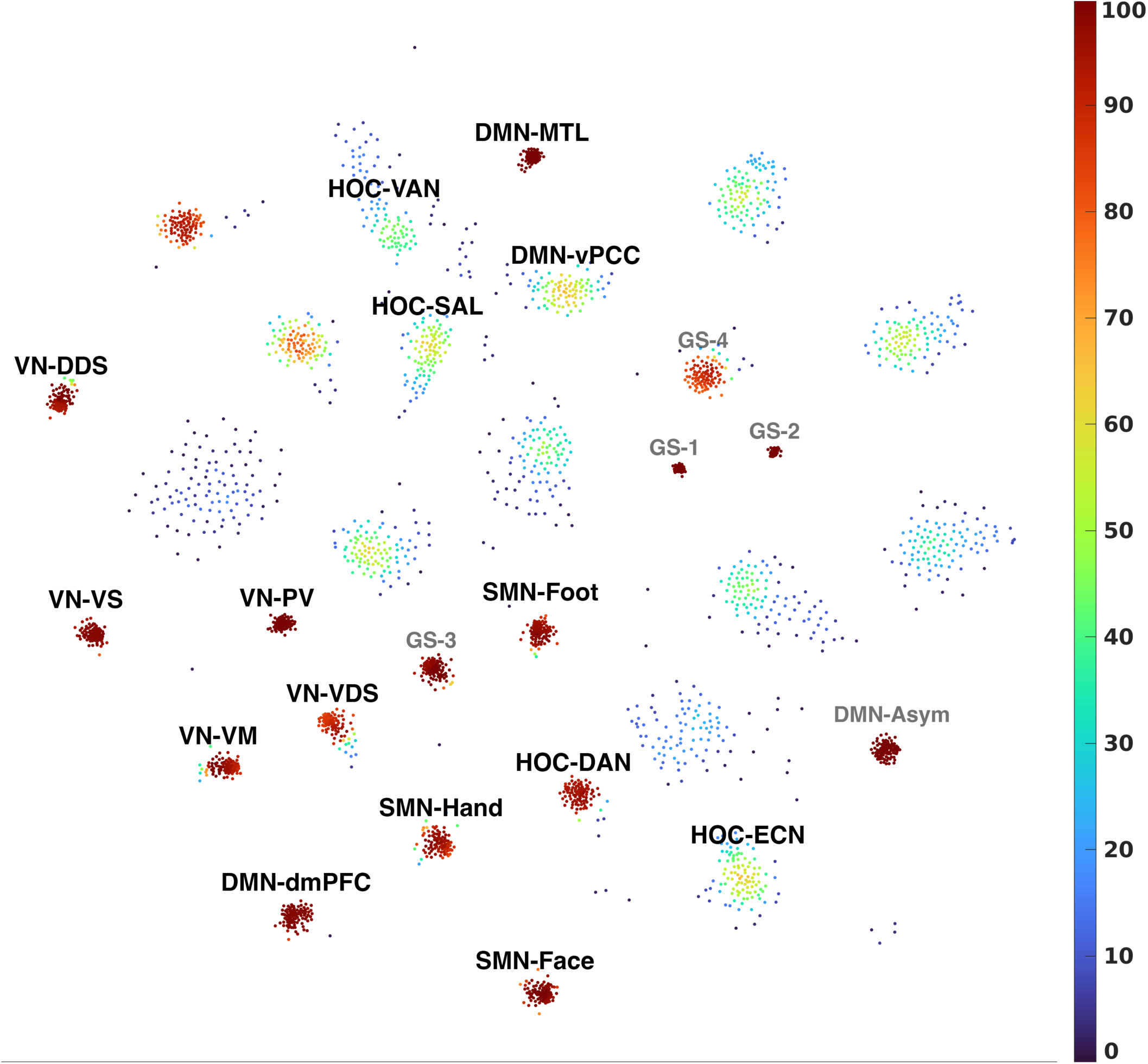
CCA-embedded spatial maps identical to Fig. S1 except for the color coding. The color bar represent the reproducibility scores, which is measured by the count of the presence of a component across all 100 bootstrap runs (a score of 100 indicates this component is found in the result from every single runs, hence highly reproducible). See Fig. S1 for other notations.

**Fig. S4.**
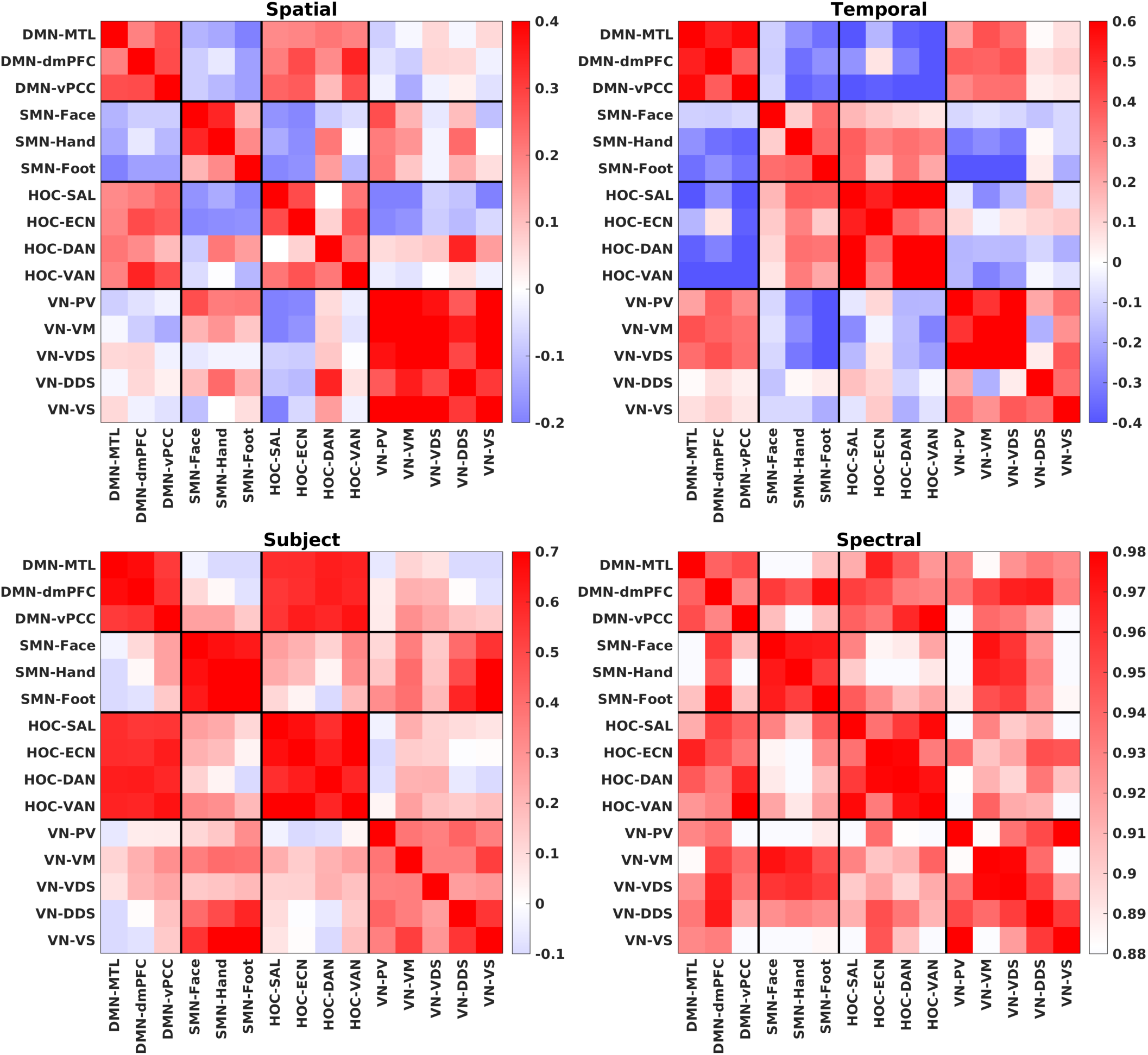
Correlation matrices among the 15 brain subnetworks. The four matrices show the correlation of their spatial maps, time series, subject modes, and spectra computed using Welch method on the time series, respectively. Red indicates positive correlations and blue indicates anti-correlations.

**Table S5.** Acronyms/Abbreviations.

|  |  |  |  |
| --- | --- | --- | --- |
| Cortical structures | (d)ACC – (dorsal) Anterior Cingulate Cortex<br>(a/dp/p)AG – (anterior/dorsal posterior/posterior) Angular Gyrus<br>cas – callosal sulcus<br>FEF – Frontal Eye Field<br>ICG – Isthmus of Cingulate Gyrus<br>IFG – Inferior Frontal Gyrus<br>ifs – inferior frontal sulcus<br>IPL – Inferior Parietal Lobule<br>(p)ITG – (posterior) Inferior Temporal Gyrus<br>(p)its – (posterior) inferior temporal sulcus<br>(a/p)MTG – (anterior/posterior) Middle Temporal Gyrus<br>(d)PCC – (dorsal) Posterior Cingulate Cortex<br>(p)PCu – (posterior) Precuneus<br>(am/dl/dm/vm)PFC – (anterior medial/dorsal lateral/dorsal medial/ventral medial) Prefrontal Cortex<br>PHG – Parahippocampal Gyrus<br>PoC – Postcentral<br>pos – parieto-occipital sulcus<br>PrC – Precentral<br>SEF – Supplementary Eye Field<br>(p)SFG – (posterior) Superior Frontal Gyrus<br>sfs – superior frontal sulcus<br>(a/p)SMG – (anterior/posterior) Supramarginal gyrus<br>SPG – Superior Parietal Gyrus<br>SPL – Superior Parietal Lobule<br>sps - subparietal sulcus<br>(p)STG – (posterior) Superior Temporal Gyrus<br>TP – Temporal Pole<br>TPJ – Temporal Parietal Junction | Thalamic nuclei | AV – Anteroventral<br>CeM – Central Medial<br>CL – Central Lateral<br>CM – Centromedian<br>LD – Lateral Dorsal<br>LGN – Lateral Geniculate Nucleus<br>LP – Lateral Posterior<br>LSg – Limitans (Suprageniculata)<br>MDl – Medial Dorsal lateral<br>MDm – Medial Dorsal medial<br>MGN – Medial Geniculate Nucleus<br>MVRe – Reuniens (Medial Ventral)<br>Pc – Paracentral<br>Pf – Parafascicular<br>PuA – Pulvinar Anterior<br>PuI – Pulvinar Inferior<br>PuL – Pulvinar Lateral<br>PuM – Pulvinar Medial<br>VA – Ventral Anterior<br>VAmc – Ventral Anterior Magnocellular<br>VL a – Ventral Lateral anterior<br>VLp – Ventral Lateral posterior<br>VM – Ventromedial<br>VPL – Ventral Posterolateral |
| Brainstem nuclei | DR – Dorsal Raphe<br>MnR – Median Raphe<br>PAG – Periaqueductal Grey<br>VTA – Ventral Tegmental Area<br>LC – Locus Coeruleus<br>LDTg – Laterodorsal Tegmental<br>mRt – Midbrain Reticular Formation<br>PBC – Parabrachial Complex<br>PnO – Pontis Oral<br>PTg – Pedunculotegmental | Basal forebrain / hypothalamic nuclei | AHA – Anterior Hypothalamic Area<br>AN – Arcuate Nucleus<br>BNST – Bed nucleus of the stria terminalis<br>DP – Dorsal periventricular nucleus<br>DM – Dorsomedial Hypothalamic<br>LH – Lateral Hypothalamus<br>MM – Mammillary bodies<br>MPO – Medial Preoptic<br>NBM – Nucleus Basalis of Meynert<br>Pa – Paraventricular<br>Pe – Periventricular Hypothalamic<br>PH – Posterior Hypothalamic<br>RN – Red nucleus<br>SC – Suprachiasmatic nucleus<br>SN – Substantia nigra<br>SO – Supraoptic Hypothalamic<br>STN – Subthalamic nucleus<br>TM – Tuberomammillary nucleus<br>VMH – Ventromedial Hypothalamic<br>ZI – Zona incerta |
| Others | SCP – superior cerebellar peduncle |  |  |

